# A mechanistic model of the spatial interaction between blue cones and blue cone bipolar cells in macaque retina

**DOI:** 10.1101/129643

**Authors:** Andreas V. Lønborg, Stephen J. Eglen

## Abstract

The spatial positions of blue cones (BC) and blue cone bipolars (BCBP) are positively correlated in macaque retina (Kouyama & Marshak, 1997): BCs, located in the outer nuclear layer (ONL), and BCBPs, located in the inner nuclear layer (INL), tend to be located close to each other in the lateral dimension. They also form separate homotypic mosaics and, finally, most BCBPs contact only a single BC pedicle. We present a mechanistic model of the BC-BCBP interaction to account for the development of these two mosaics and their connections. It assumes that BCs are immobile and that BCBPs can migrate within the INL due to a compromise between a dendritic string force and an intrinsic INL friction force. Model parameters were selected to optimise the fit with observed fields (Kouyama & Marshak, 1997). Simulated density recovery profiles (DRPs) closely mimic the observed DRPs. In particular, for each of the five retinal fields studied, the simulated DRP for the interaction between BCs and BCBPs has a peak at distances less than around 20 μm and a small dip at distances up to the maximal lateral dendritic length (∼ 44 μm), matching the profiles of the real data. We conclude that our mechanistic model is a candidate for the unknown mechanism that drives the observed interaction between BCs and BCBPs in macaque retina.

## Abbreviations

BC: Blue Cone
BCBP: Blue Cone Bipolar
DRP: Density Recovery Profile
INL: Inner Nuclear Layer
ONL: Outer Nuclear Layer
SD: Standard Deviation

## Introduction

Many types of retinal neurons are organised in regularly distributed patterns, called *retinal mosaics*. Pioneering quantitative work on retinal mosaics was initiated by Wässle & Riemann (1978), who found that a selection of retinal cell types (including cone photoreceptors, horizontal cells and ganglion cells) were organised in mosaic patterns. To generate such patterns, it is thought that neurons must interact with each other locally, since a global blueprint of the mosaic would require more information than could be stored in the progenitor cells. Since Wässle and Riemann reported their pioneering results, a large number of other empirical studies have identified and quantified mosaics for various cell types and in various species, reviewed e.g. by Cook & Chalupa (2000).

In general, the types of developmental interactions between arrays of cells can be grouped into three categories (Kouyama & Marshak, 1997): (i) interactions between cells of the same type, (ii) between cells of different type but in the same layer and (iii) between cells of different type in different layers. The first type of interaction is called *homotypic* and the latter two are called *heterotypic*.

Most studies have focused on homotypic interactions but several studies have searched for heterotypic interactions between different cell types within the same layer (Wikler & Rakic, 1991; Curcio et al., 1991; Bumsted et al., 1997; Zhan & Troy, 2000; Eglen et al., 2005; Eglen & Wong, 2008) and between cell types in different layers (Kouyama & Marshak, 1997; Luo et al., 1999; Ahnelt et al., 2000; Rockhill et al., 2000; Eglen et al., 2003; Mack, 2007). The general conclusion is that heterotypic interactions are much less prominent than homotypic interactions. There are, however, notable exceptions (Kouyama & Marshak, 1997; Ahnelt et al., 2000).

In the macaque retina, BCs in the ONL and BCBPs (a subclass of bipolar cells which connect exclusively to BCs) in the INL are spatially positively correlated (Kouyama & Marshak, 1997). So far, this interaction has not been explained satisfactorily. Our study seeks to fill this gap through the development of a mechanistic model for the interaction between BCs and BCBPs.

When developing a model, one needs to decide on the presumptive mechanism(s) that drive pattern formation. Over the years, various competing explanations of retinal mosaics have been suggested; for reviews see Cook & Chalupa (2000); Eglen & Galli-Resta (2006). In brief, the suggested mechanisms are (i) lateral inhibition of cell fate (ii) cell death (iii) lateral migration and (iv) dendritic interactions. The present study argues in favour of the latter two by showing that in the case of the macaque retina, BCBP migration induced by dendritic interactions between BCs and BCBPs is sufficient to reproduce the observed positive spatial correlation.

Another characteristic of the model developed here is that it is in accordance with the following biological facts about BCs and BCBPs. First, BCs are packed densely in the ONL (Bumsted et al., 1997) and their axons and pedicles have been shown to be immobile (Ahnelt & Kolb, 2000). Hence, BCs have been assumed to be fixed. Second, BCBPs tend to contact fewer cones than diffuse bipolars and a large proportion of them make only one connection (Kouyama & Marshak, 1992; Herr et al., 2003).

Assuming that the dendrites exert a string-like force on the BC pedicles and the BCBPs, we speculated that the observed positive correlation could primarily be caused by migration of those BCBPs with only one dendritic connection to BC pedicles. The mechanism is remarkably simple: if there is a force between two bodies and the two bodies are confined to parallel but separate layers, then this force will tend to reposition the bodies such that the lateral distance between the bodies is reduced. By lateral distance we mean the distance as measured when cells are projected onto a plane parallel with the retinal layers. Lateral distance therefore disregards the radial displacement of cells.

In the retina, the BCBPs cannot migrate freely, so the model should not imply perfect radial alignment of BCs and single dendrite BCBPs. The constraints on BCBP migration are assumed to be equivalent to a friction force in the INL, and we have therefore incorporated friction in the model.

According to the ideas sketched out above, we have developed a mechanistic string and friction force model for the interaction between BCs and BCBPs in macaque retina. Using this model, patterns of BCs and BCBPs were generated using dimensions and cell numbers identical to the observations by Kouyama & Marshak (1997). The results show that the devised model is capable of reproducing the empirically observed spatial correlation pattern between BCs and BCBPs.

## Methods

### Data sets

Real data fields were provided by Kouyama & Marshak (1997). The data consist of the spatial positions of BCs and BCBPs in five midperipheral retinal fields (named here F1–F5) from the same monkey. The data were obtained using a double labelling procedure as described by Kouyama & Marshak (1992). The fields vary in BC density (274–470 mm^*-*2^) and BCBP density (544–940 mm^*-*2^), with F4 having the highest BC density and F1 having the highest BCBP density. Details of the fields are provided in Table 1, see also Table 1 in Kouyama & Marshak (1997). When it comes to testing the mechanistic model, it is a clear advantage that there is variation among the five fields, since this implies that the model will be applied to five different scenarios.

**Table 1:**
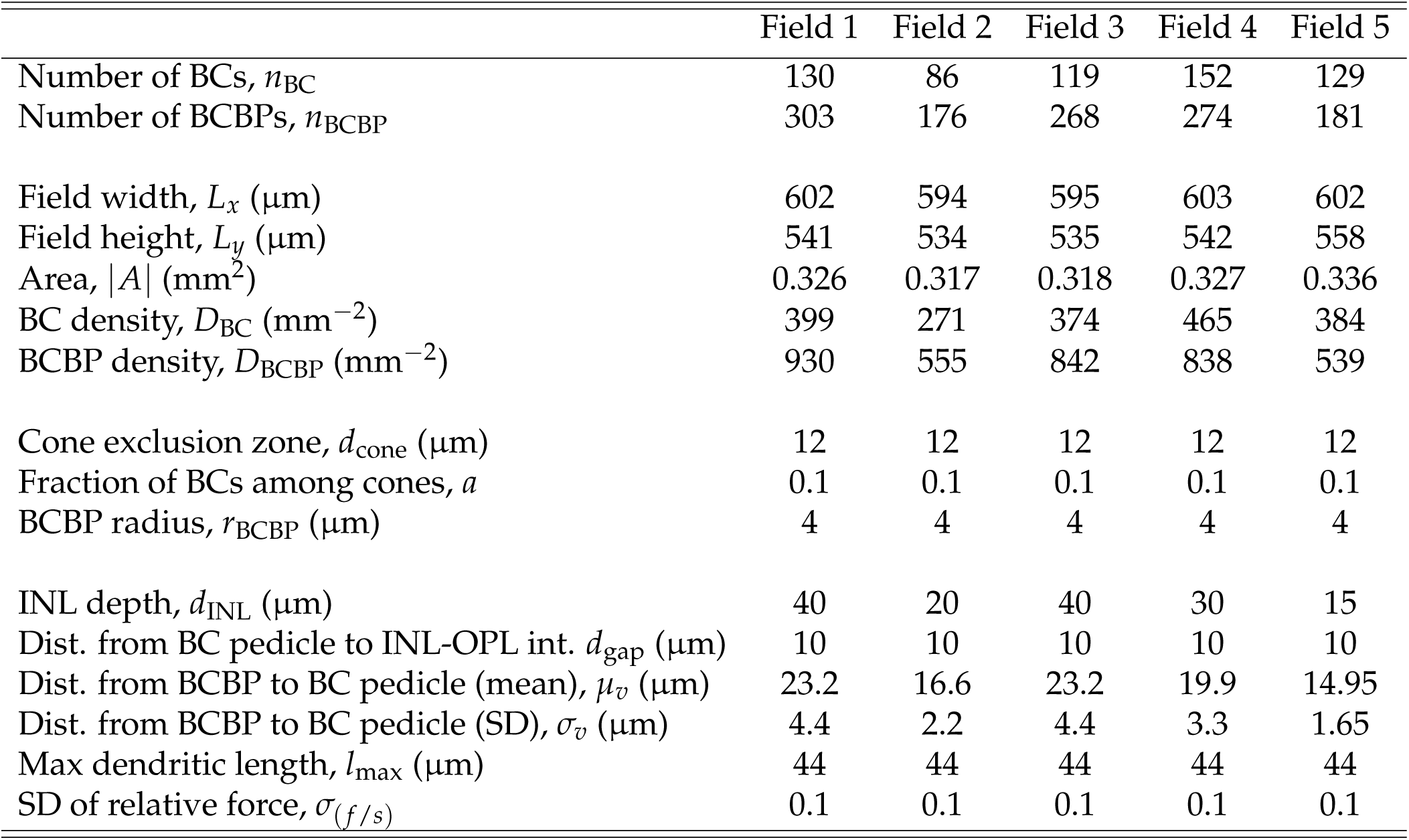
Values of fixed parameters of the model. Where possible, these parameters have been estimated from the literature or measured from the real data sets, and were not changed during model fitting.

|  | Field 1 | Field 2 | Field 3 | Field 4 | Field 5 |
| --- | --- | --- | --- | --- | --- |
| Number of BCs, $n_{\text{BC}}$ | 130 | 86 | 119 | 152 | 129 |
| Number of BCBPs, $n_{\text{BCBP}}$ | 303 | 176 | 268 | 274 | 181 |
| Field width, $L_x$ ($\mu\text{m}$) | 602 | 594 | 595 | 603 | 602 |
| Field height, $L_y$ ($\mu\text{m}$) | 541 | 534 | 535 | 542 | 558 |
| Area, $\vert A\vert $ ($\text{mm}^2$) | 0.326 | 0.317 | 0.318 | 0.327 | 0.336 |
| BC density, $D_{\text{BC}}$ ($\text{mm}^{-2}$) | 399 | 271 | 374 | 465 | 384 |
| BCBP density, $D_{\text{BCBP}}$ ($\text{mm}^{-2}$) | 930 | 555 | 842 | 838 | 539 |
| Cone exclusion zone, $d_{\text{cone}}$ ($\mu\text{m}$) | 12 | 12 | 12 | 12 | 12 |
| Fraction of BCs among cones, $a$ | 0.1 | 0.1 | 0.1 | 0.1 | 0.1 |
| BCBP radius, $r_{\text{BCBP}}$ ($\mu\text{m}$) | 4 | 4 | 4 | 4 | 4 |
| INL depth, $d_{\text{INL}}$ ($\mu\text{m}$) | 40 | 20 | 40 | 30 | 15 |
| Dist. from BC pedicle to INL-OPL int. $d_{\text{gap}}$ ($\mu\text{m}$) | 10 | 10 | 10 | 10 | 10 |
| Dist. from BCBP to BC pedicle (mean), $\mu_v$ ($\mu\text{m}$) | 23.2 | 16.6 | 23.2 | 19.9 | 14.95 |
| Dist. from BCBP to BC pedicle (SD), $\sigma_v$ ($\mu\text{m}$) | 4.4 | 2.2 | 4.4 | 3.3 | 1.65 |
| Max dendritic length, $l_{\text{max}}$ ($\mu\text{m}$) | 44 | 44 | 44 | 44 | 44 |
| SD of relative force, $\sigma_{(f/s)}$ | 0.1 | 0.1 | 0.1 | 0.1 | 0.1 |

### Modelling blue cones and blue cone bipolars

The model developed in this paper is divided into five steps, which correspond to five postulated phases in the development of the BC and BCBP arrays. (i) Generation of undifferentiated cones, (ii) differentiation of BCs, (iii) generation of BCBPs, (iv) formation of synaptic connections between BCBP dendrites and BC pedicles and (v) lateral migration of BCBPs due to a compromise between the dendritic string force and the INL friction force. Note that nothing will be assumed about the relative timing of phase (i)/(ii) and (iii), although it is likely that cones differentiate before bipolar cells (Rapaport, 2006).

The description below of the five steps corresponds to the simulation of one generic version of these fields. The size of the simulated fields and the number of BCs and BCBPs in each of these mimic the real data. This implies that the simulated field densities are almost identical to the real densities. The minor differences are caused by a slightly different definition of field size compared with Kouyama & Marshak (1997). Finally, steps 2–5 of the model were repeated *N* = 99 times in order to reduce the uncertainty in the results caused by random sampling. We have divided the parameters of the model into two; those that are fixed by the observed data (Table 1), and those that are free (Table 2).

**Table 2:**
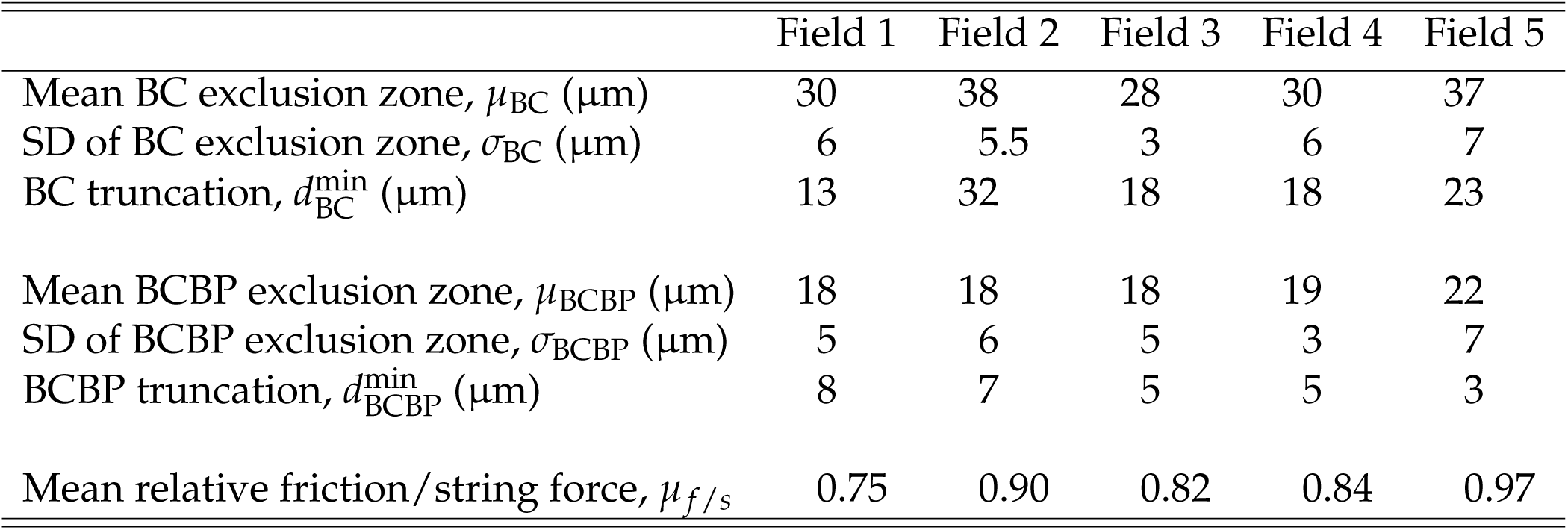
Free parameters for each field. These parameters were found by trial-and-error exploration of the model.

### Step 1, generation of cones

A set of *n*_*BC*_/*a* undifferentiated cones were generated by drawing points from a bivariate uniform distribution on the area of the field. Cones were subjected to a *d*_cone_ = 12 μm exclusion zone. This avoids physical overlap between BCs, which have a diameter of around 10 μm (Kouyama & Marshak, 1992) and the pedicles of which lie approximately in the same plane (Kolb et al., 1997). Since cones are relatively densely packed (Bumsted et al., 1997) there is only moderate scope for randomness, so the bias introduced by reusing the same set of cones in each of the *N* = 99 simulations is likely to be negligible. Thus, for simplicity, the same set of undifferentiated cones was used in all *N* = 99 simulations. From this point onwards the cones do not move. This assumption is supported by the observation that the rods and cones in the photoreceptor layer are densely packed (Bumsted et al., 1997) and also by a study which shows that the axons and pedicles of BCs do not move (Ahnelt et al., 2000).

### Step 2, differentiation of blue cones

In mammals, BC opsin is expressed prior to green and red cone opsin (macaque (Bumsted et al., 1997), human (Xiao & Hendrickson, 2000), rabbit and rat (Szél et al., 1994), gerbil and mouse (Szél et al., 1993)). Therefore, BC differentiation was modelled as a process where a fraction of undifferentiated cones differentiate into BCs. According to Bumsted et al. (1997) a fraction *a* = 10 % of cones in the macaque are BCs. In our model, this fraction of spatially fixed cones thus differentiates into *n*_BC_ BC BCs under the constraint that each BC has an individual exclusion zone drawn from a truncated Gaussian with mean *μ*_BC_, standard deviation (SD) *σ* _BC_ and truncated below at 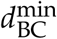. At this stage, a lattice of cones, some of which are BCs, has been generated. An example lattice is shown in Figure 1.

**Figure 1:**
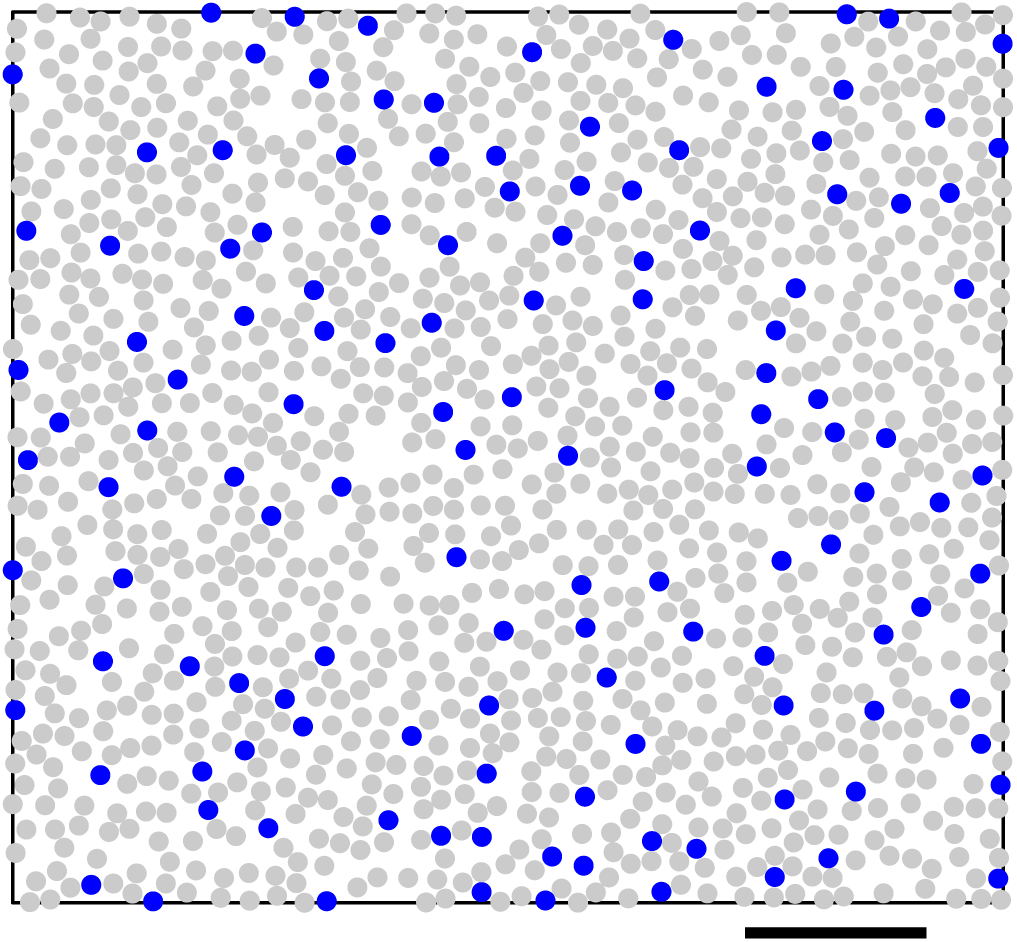
Example of BC differentiation (filled blue circles) from undifferentiated cones (filled light grey circles) for a simulation of field 1. Each BC has an individual exclu-sion zone. Circle diameter matches approximate size of BC (cone diameter is roughly 10 μm). Scale bar is 100 μm.

### Step 3, generation of blue cone bipolars

The *n*_BCBP_ bipolar cells were given an initial (*x*, *y*) location in the same way as the undifferentiated cones (step 1), with the only exception being that each BCBP was assigned an individual exclusion zone drawn from a truncated Gaussian with mean BCBP *μ*_BCBP_, SD *σ* _BCBP_ and truncated below at 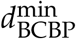. As the INL is much deeper than the diameter of a single BCBP, BCBPs were also given a depth within the INL according to observations in Kouyama & Marshak (1992). Specifically, depths were drawn from a Gaussian distribution with mean 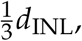, where *d*_INL_ is the depth of the INL, and SD *σ* _*v*_ = 0.11*d*_INL_. Values for *d*_INL_ were estimated by using the information about *d*_INL_ at various distances from the fovea in Figure 2 of Kouyama & Marshak (1992) and comparing with the distances from the fovea of the five fields (given in Table 1 of Kouyama & Marshak, 1997). The distance between the INL-OPL interface and the BC pedicles, *d*_gap_, is approximately 10 μm (estimated from Figure 2 of Kouyama & Marshak, 1992); hence, the vertical distance *d*_*v*,*i*_ between the plane of BC pedicles and BCBP_*i*_ was drawn from a Gaussian distribution with mean 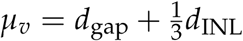 and SD *σ* _*v*_ = 0.11 *d*_INL_. See Figure 2 for a diagram indicating these depth variables. The space between the INL-OPL interface and the BC pedicles is void of somata.

**Figure 2:**
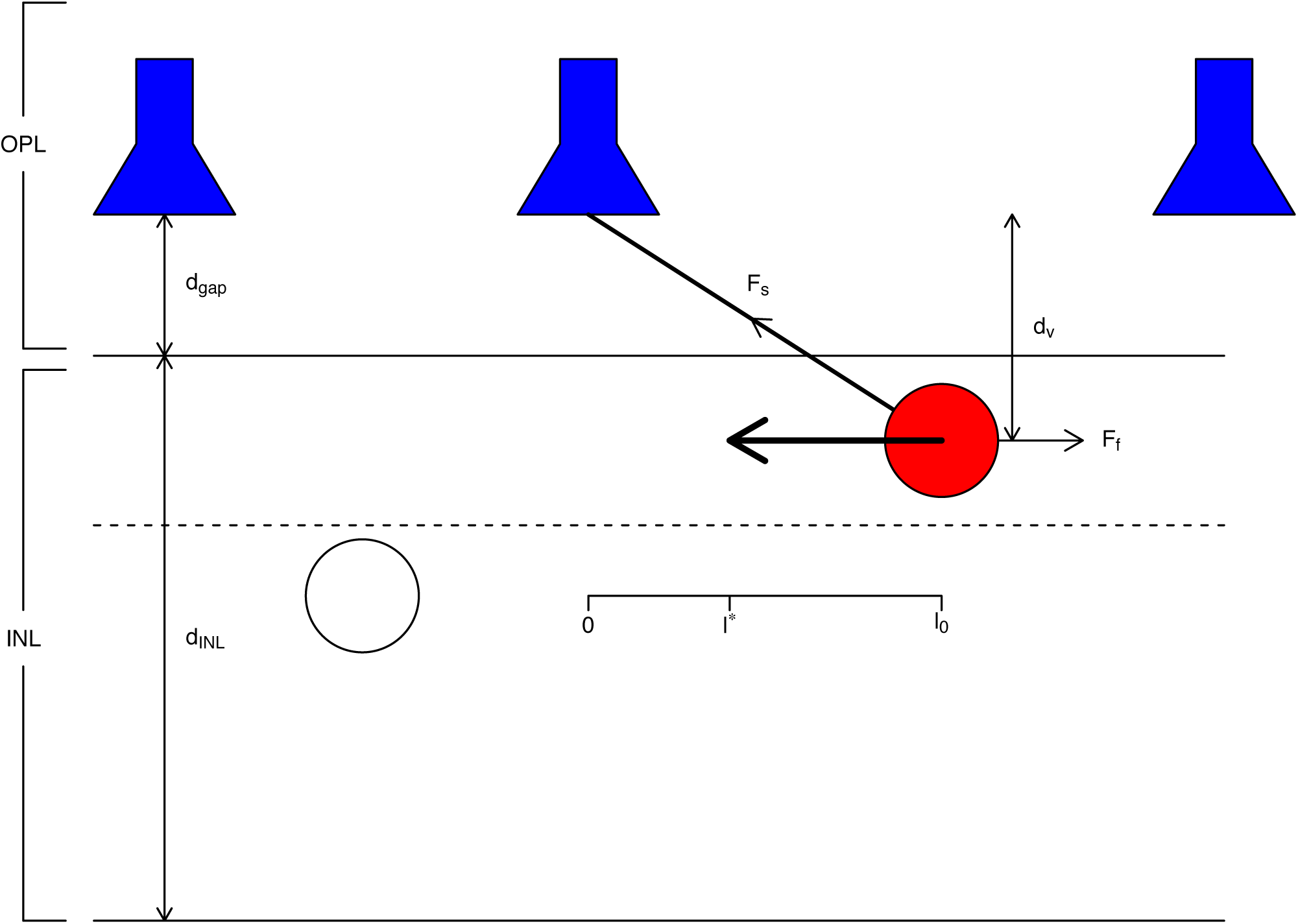
Diagram of the mechanistic model. BC pedicles (top, shown in blue) are fixed in the OPL, whereas BCBPs which are less than 3 *r*_BCBP_ = 12 μm (threshold shown by dotted line) from the INL-OPL interface can migrate laterally. BCBPs which are located deeper in the INL (e.g. the open circle) are assumed to be immobile because their dendrites will potentially not contact a BC pedicle in a straight line because of other BCBPs blocking the way. The filled red BCBP can move and is connected to the central BC pedicle. The BCBP is under the influence of a dendritic string force **F**_*s*_ and an opposing friction force **F** _*f*_. As BCBPs are constrained to move only laterally, the BCBP migrates from an initial position *l*_0_ to postmigrational position *l*^*^ (indicated by inset ruler). The thick arrow indicates the migration. Relevant depth parameters (*d*_gap_, *d*_INL_ and *d*_*v*_) are indicated.

### Step 4, formation of synaptic contacts between blue cones and blue cone bipolars

Information about BCBP dendrites was used in the modelling of the synaptic contacts between BCs and BCBPs. Kouyama & Marshak (1992) reported that, on average, a macaque BCBP makes 1.2 synaptic contacts with BCs (this number is called the *convergence*), whereas BCs make 1.8 contacts with BCBPs (this number is called the *divergence*). Also, almost 70 % of all BCBPs make synaptic connections to only one BC. Less than 30 % make more than one connection. In our work, we assumed that no BCBP can make more than one contact. Furthermore, following the observation in Kouyama & Marshak (1992) that no dendrite is longer than 50 μm, we assumed that BCBPs, with a lateral distance of more than *l*_max_ = 44 μm from the nearest BC, do not form contacts. (This assumes that the mean depth of a BCBP cell is 23.3 μm from the max BC pedicles, see step 3, and thus 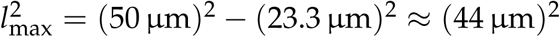 Under these assumptions, each BCBP was connected to the nearest BC. Finally, and also following an observation in Kouyama & Marshak (1992), BCs were restricted to have a maximum of four connections to BCBPs. (If during this stage a BC had more than four connections, connections were removed randomly until it had four connections.)

### Step 5, lateral migration of blue cone bipolars

At this stage, BCBPs have formed connections to BCs and now face two opposing forces: (i) the dendritic string force and (ii) a friction force which resists migration in the INL. BCBPs are assumed to migrate laterally as a compromise between these two forces. The assumed physical situation is sketched in Figure 2.

As BCBPs are located at various depths within the INL, we assumed that if a BCBP is located in a way such that another BCBP can potentially be positioned radially towards the INL-OPL interface relative to this BCBP, then it cannot move. Therefore, a BCBP cell can move laterally only if it is within a distance of 3 *r*_BCBP_ = 12 μm from the the INL-OPL interface. (The radius of a BCBP soma is estimated to be *r*_BCBP_ = 4 μm (Kouyama & Marshak, 1992)). This implies that in the case of *d*_INL_ = 40 μm, 38 % of BCBPs have the potential to migrate (since *P*(*X <* 12)= 0.38, where 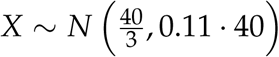. By restricting the number of BCBPs which can migrate, the output correlation between BCs and BCBPs is effectively reduced.

Assuming that BCBP_*i*_ can migrate laterally (i.e. *d*_*v*,*i*_ *< d*_gap_ + 3 *· r*_BCBP_), the amount of movement is a trade-off between the dendritic string force **F**_*s*,*i*_ (which depends on the vertical distance *d*_*v*,*i*_) and the friction force **F** _*f*,*i*_. We assumed there is no difference between static and dynamic friction forces, and that the friction force **F** _*f*,*i*_ on BCBP_*i*_ is velocity independent and directed against the direction of migration. Finally, it was assumed that dendrites adapt during migration, such that the dendrites are always taut.

The postmigrational equilibrium position of a BCBP can be derived by writing the equation of motion for the system and finding the steady state (where acceleration and velocity are zero). The derivation is given in the Supplementary material, section 1; the result is:

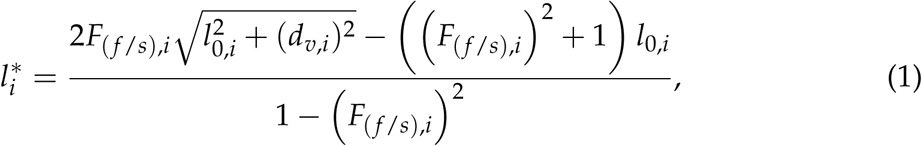

where 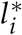 denotes the postmigrational lateral position of BCBP_*i*_ relative to the BC pedicle, *l*_0,*i*_ denotes the initial lateral distance between BCBP_*i*_ and the BC pedicle it connects to, *d*_*v*,*i*_ denotes the vertical distance, and *F*_(*f*/*s*),*i*_ *≡ F*_*f*,*i*_/*F*_*s*,*i*_ denotes the relative friction-to-string force. The postmigrational position depends only on the ratio between the friction and the string force *F*_(*f*/*s*)_ and not on their absolute sizes. Given that we have no *a priori* information about the values of these forces, it is an advantage that we have to deal with only one free parameter since it makes the model more parsimonious.

In the simulations, each BCBP was assigned a relative force 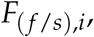, drawn from a Gaussian distribution with mean *μ* _*f*/*s*_ and SD *s*_*f*/*s*_ = 0.1. Once 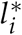 was calculated for each BCBP_*i*_ that could migrate, the cells were moved laterally to their new position. This ignores the possibility that BCBPs might collide with each other during migration. To alleviate this, once BCBPs were moved, any pair of BCBPs that were too close to each other were found and the their migration was reversed in small steps until the somata no longer overlapped. This approach to simulating migration was chosen as it was more efficient than developing and running a time-resolved simulation to track the dynamic movements of each cell during migration.

### Quantitative model output

To assess whether the model can generate the cross-correlations observed in real data, we used density recovery profiles (Rodieck, 1991), as also used by Kouyama & Marshak (1997). The DRP provides an estimate of the density of cells of type A as a function of the distance from a cell of type B. To emphasise when we are correlating two different populations, we call the result a ‘cross DRP’; when a population of cells is correlated against itself, we call the result just ‘DRP’. Regularity of the individual mosaics was also assessed using the nearest-neighbour regularity index (Wässle & Riemann, 1978). We also calculated other properties of the resulting patterns, which serve as predictions for future work. The adjustable parameters of the model were manually selected to give the best fit. Computational modelling and analysis were performed using the R programming environment (R Development Core Team, 2008). Code and data are available upon request.

## Results

Figure 3 shows a simulated field and the corresponding real data field, using the parameters listed in Table 2. Figure 3 demonstrates the significant difference in BC and BCBP densities (see also Table 1). Further, the mosaic structure of both BCs and BCBPs is clearly visible in both the real data field and the simulated data field. Given that only one simulated example is shown, we do not expect it to be optimal, and although it looks comparable, quantitative methods are required to further compare model and data.

**Figure 3:**
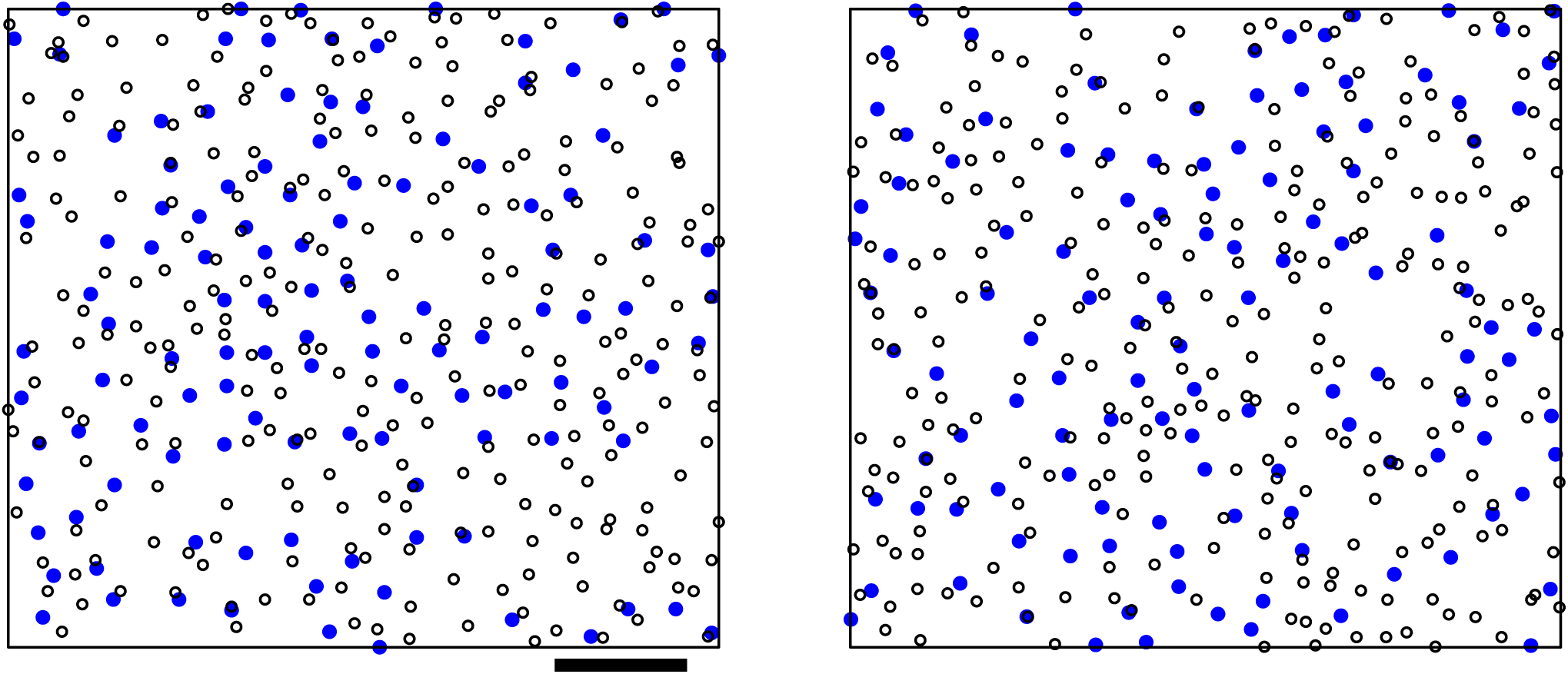
Comparison of a real data field (field 3, left) and a corresponding simulated field (right). Filled blue circles indicate BCs and open circles indicate BCBPs. In both fields, both BCs and BCBPs are arranged in mosaics; also the simulated field looks similar to the real data field. The relative sizes of cells and field are to scale. Scale bar is 100 μm.

To allow for a thorough comparison of model with observed data, we ran the model *N* = 99 times so that we could assess the average performance of the model, which should be more representative than looking at any one simulation. In particular, results from simulations are summarised, for a particular measure, as the mean *±* 1.96 SDs across simulations, to indicate the 95% confidence interval of that measure (under normality assumptions).

As a first step to comparing real and simulated data, we computed the traditional measure of mosaic regularity, the regularity index (Wässle & Riemann, 1978), individually for the BC mosaic and the BCBP mosaic (Figure 4). For all but one mosaic, the observed regularity index falls within the error bars of the simulations. This suggests that the model is capable of reproducing the statistics of individual mosaics, which is a key prerequisite before we can investigate the key component of the model, the nature of the cross-correlation.

**Figure 4:**
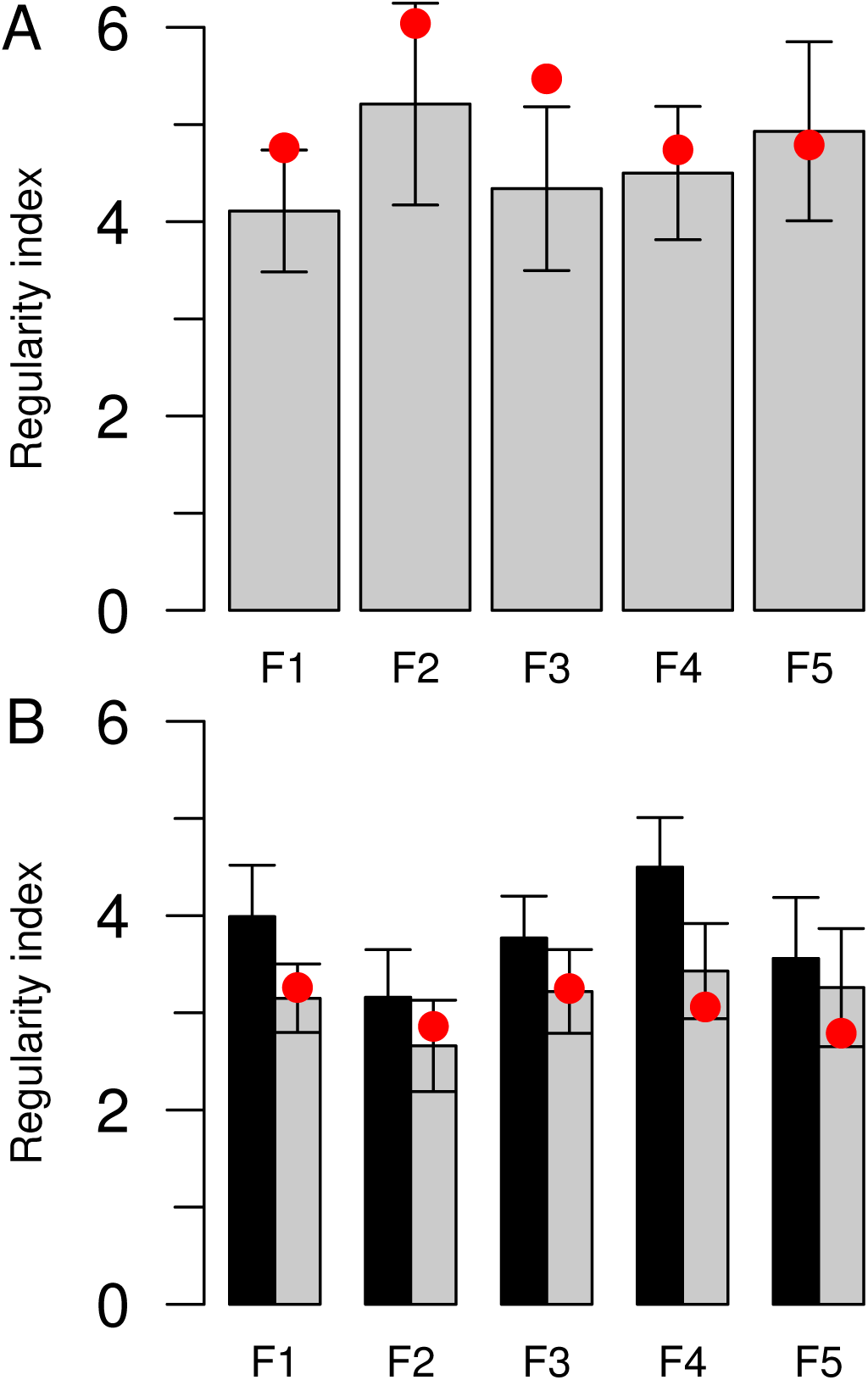
Nearest-neighbour regularity index of each mosaic. Bars show the mean across simulations, and error bars denote the mean *±*1.96 times the SD across simulations (*N* = 99 simulations). The filled dots indicate the observed regularity index for that field. (A) Regularity of BC mosaic. (B) Regularity of BCBP mosaic before (black bars) and after migration (grey bars).

Figure 5 gives an insight into the dynamics of the model, from which three observations can be made. (i) Most BCBPs do not migrate, (ii) those that migrate do not tend to migrate very far and (iii) migration implies that BCBPs move closer to BCs. These output characteristics are not surprising given the nature of the model, but they constitute predictions which could potentially be tested against empirical observations. Therefore it is useful to make them more quantitative.

**Figure 5:**
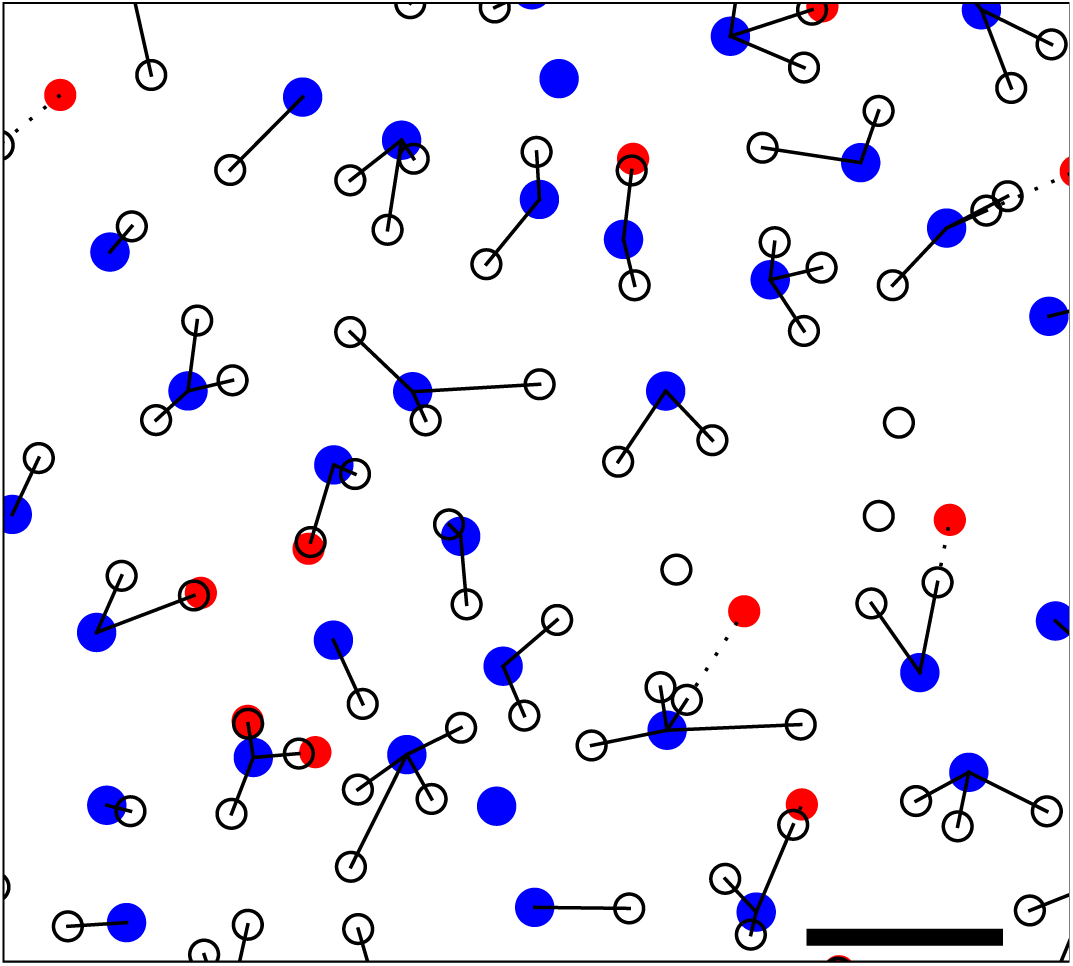
BCBP migration and dendritic contacts with BCs for one simulation of field 3 (central 1/4 of field shown). BCs are indicated as filled blue circles, BCBPs as open circles and the initial positions of those BCBPs that migrate are indicated as red circles. Postmigrational dendrites are illustrated with solid lines and BCBP migration is illustrated with dotted lines. Most BCBPs do not migrate and those which do move closer to the nearest BC. The relative sizes of cells and field are to scale. Scale bar is 50 μm.

The proportion of BCBPs that migrate is illustrated in Figure 6; only 10–20 % of BCBPs migrate once BCBPs have potentially made contacts with BCs. This is a firm prediction of the model. Field 2 stands out as having the highest migration rate. The low BC density in this field implies that the dendrites tend to be longer, which in turn implies that the projected string force will be stronger, thus increasing the likelihood of migration.

**Figure 6:**
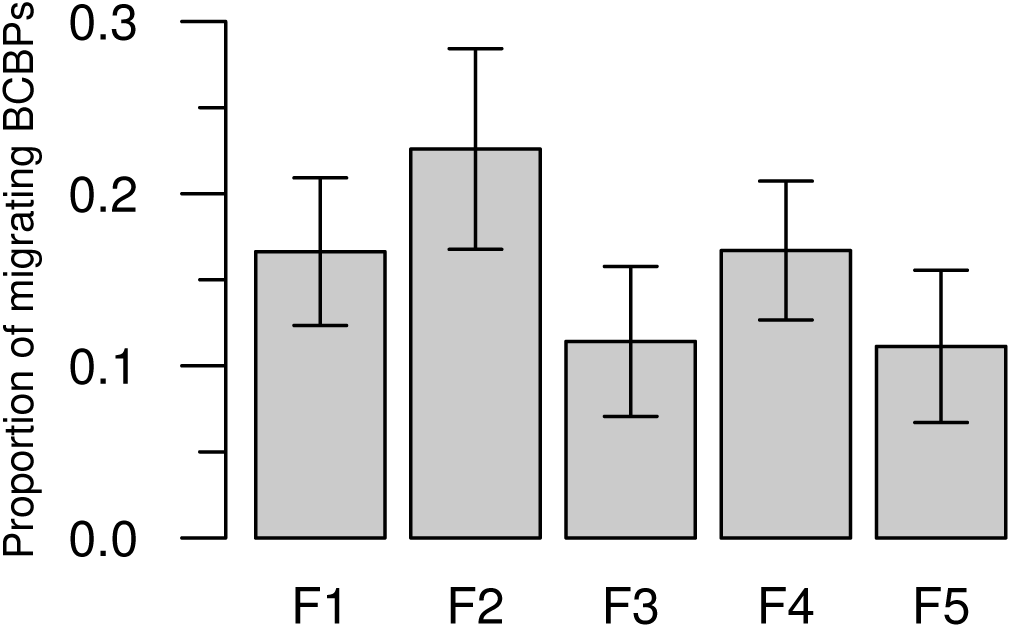
Proportion of BCBPs that migrate in each field. Bars show the mean across simulations, and error bars denote the mean *±*1.96 times the SD across simulations (*N* = 99 simulations).

Figure 7 shows the average dendritic length of all BCBPs before and after BCBP migration. The average premigration values are around 20 μm, which compares with a analytical prediction of 33 μm (see Supplementary material, section 2). The analytical calculation ignores the issue that as the distance from the BC increases, the chance that a BCBP will connect to another BC increases. Also, a few BCBPs do not make any connections, probably because they are too far from BCs. Accounting for both these factors would reduce the theoretical premigrational dendritic length, and so the value of 33 μm should be regarded as an upper bound. In this context, the average premigration distance of around 20 μm from the simulations seems reasonable.

**Figure 7:**
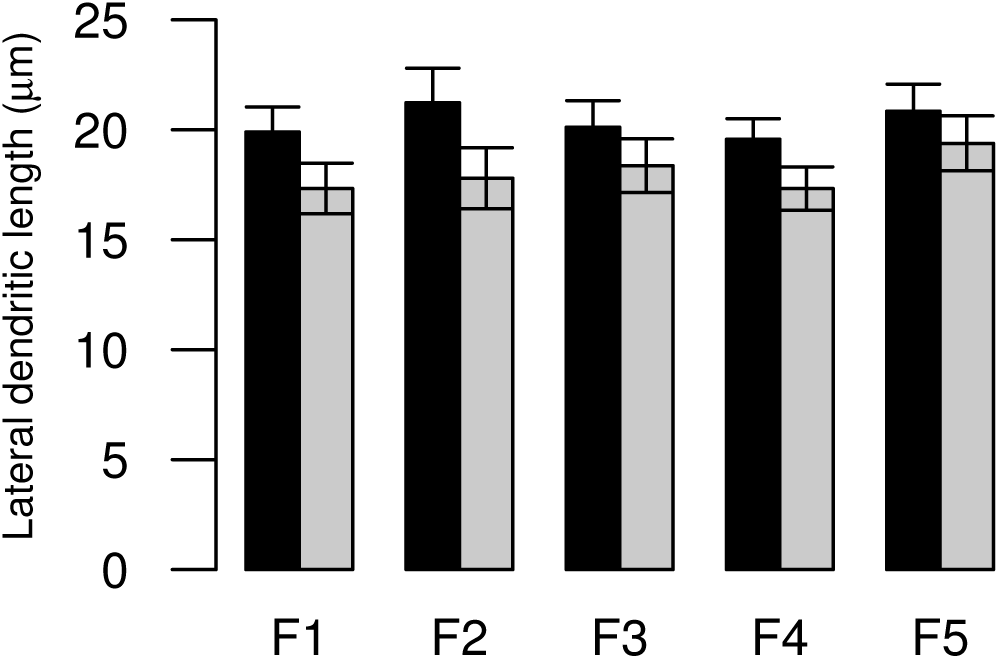
Pre-(black bars) and postmigrational (grey bars) dendritic lengths for all BCBPs for each field. Bars show the mean across simulations, and error bars denote the mean *±*1.96 times the SD across simulations (*N* = 99 simulations).

Furthermore, the premigrational lengths are largest for field 2. This is caused by the lower density of BCs in this field. To understand this, note that if there were infinitely many BCs, the dendritic lengths would be *l*_0_ = 0 for all BCBPs. Reducing the number of BCs increases the distance of a BCBP to its nearest BC. Some BCBPs become too distant from BCs and will not make a connection. This increase will stop only when there are so few (and regularly distributed) BCs that each BCBP can maximally reach only one BC. This point is not reached in the fields studied here. Hence, the lower the BC density, the larger the premigrational dendritic length.

It is tempting to reverse this argument and conclude that the fewer BCBPs the smaller *l*_0_, but this is flawed since BCs and BCBPs are not symmetric. BCBPs can grow only one dendrite which connects to a BC pedicle, whereas a BC pedicle can connect to as many as four BCBP dendrites.

The difference between preand postmigrational dendritic lengths is around 2–3 μm, but this number does not reflect true migration distances since these values are averages of all BCBPs, i.e. also including those which do not move. When considering only the BCBPs that move, migration distances average around 15 μm (Figure 8), although there is substantial variation in migration distances across simulations.

**Figure 8:**
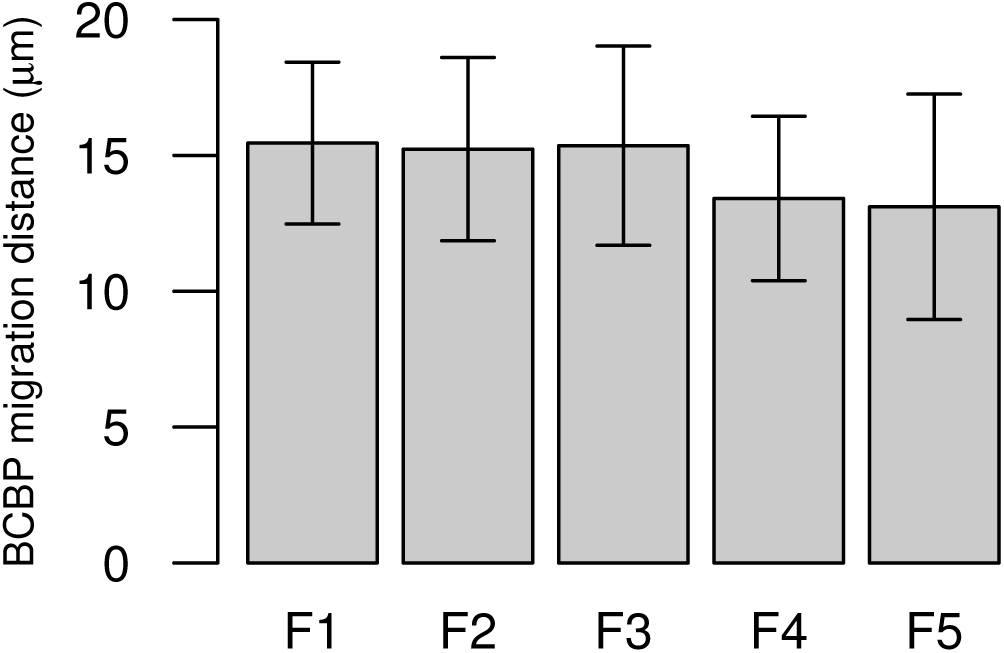
Migration distances for BCBPs in each field. Only BCBPs that migrate are considered here. Bars show the mean across simulations, and error bars denote the mean *±*1.96 times the SD across simulations (*N* = 99 simulations).

Finally, the number of synapses in the simulated model can be compared with the real number of synapses (Figure 9). The fit with real data is quite good, which is surprising given that the number of synapses is an endogenous part of the model and that we assumed that BCBPs either connect to zero or one BC, yet in reality BCBPs can connect to two BCs (Kouyama & Marshak, 1992). Hence, we would expect that the simulations generate fewer synapses than there are in the real data. Figure 9 shows that this is true only for fields 4 and 5. In the case of fields 1–3, the simulated number of synapses is larger than the observed number. This implies that the simulations of field 1–3 might have a higher capability to create BC-BCBP interactions than the real data. If the number of synapses in fields 1–3 were reduced, then the peaks in the cross DRP, see next section, would become less pronounced (given the relative friction force) and a smaller relative friction force would be needed to reproduce the sizes of the peaks observed in real data. The point of this argument is that the estimated relative friction forces are not sufficiently accurate to be interpreted as the true physical relative forces. A more detailed model is needed if such interpretation should be valid. Nonetheless, the estimated relative force provides a good first approximation.

**Figure 9:**
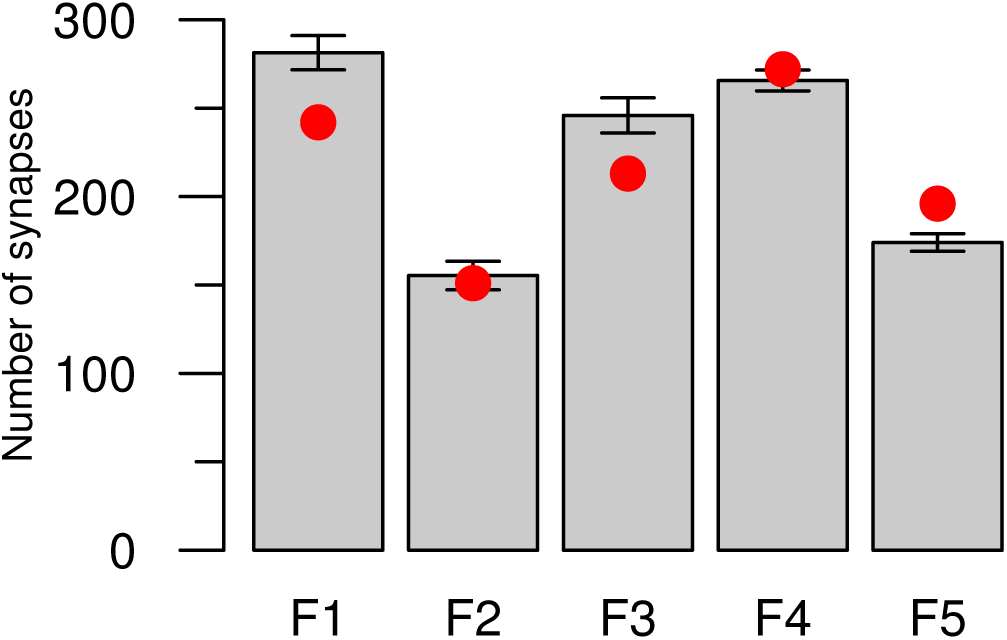
Number of synapses created in each of the five fields. The number of synapses in the real data fields is indicated by filled dots. Bars show the mean across simulations, and error bars denote the mean *±*1.96 times the SD across simulations (*N* = 99 simulations).

### Density recovery profiles

The DRP analysis is the most important analysis performed here, since this is the analysis which quantifies the correlation between BCs and BCBPs. For each field, five different DRPs were generated. One DRP for the BCs, two DRPs for the BCBPs (before and after BCBP migration) and two cross DRPs for the BC-BCBP interaction (before and after BCBP migration). For a representative field, field 2, the five DRPs are shown in Figure 10. For the other fields, only the postmigrational cross DRPs are shown (Figure 11).

**Figure 10:**
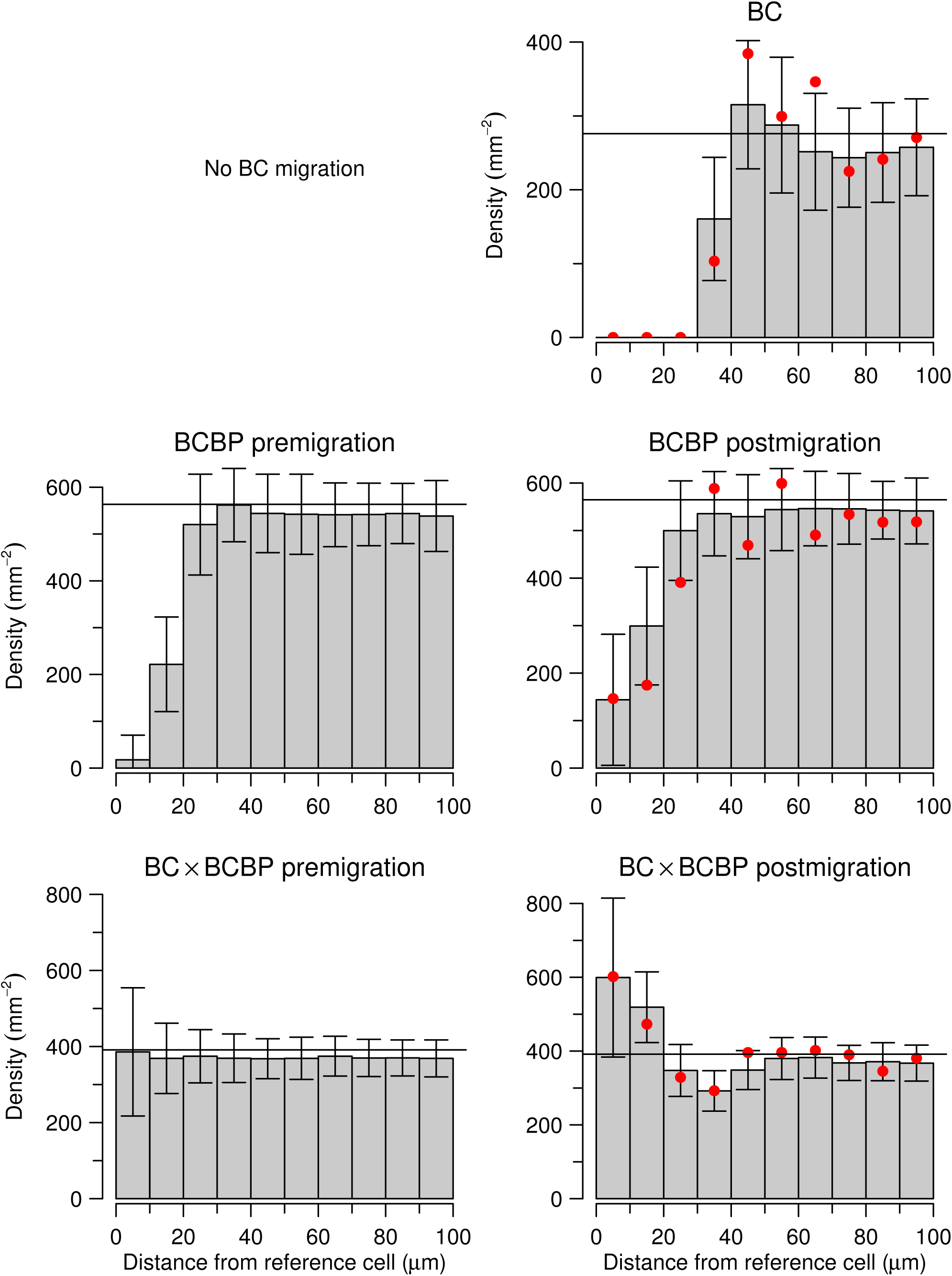
DRP results for field 2. Left column shows DRPs before migration, and right column shows DRPs after BCBP migration. Top row: DRP for BCs (as BCs do not migrate, the DRPs before and after migration are identical). Middle row: DRPs for BCBPs. Bottom row: cross DRPs for BC *×* BCBPs. Filled dots indicate real data results and horizontal lines indicate macroscopic densities. Bars show the mean across simulations, and error bars denote the mean *±*1.96 times the SD across simulations (*N* = 99 simulations).

In Figure 10, the BC DRP has a characteristic dip at small distances, which reflects the homotypic exclusion zone. At larger distances the DRP ‘recovers’ to the macroscopic density (indicated by a horizontal line). The same is the case for the premigrational BCBP DRP, with the difference that the exclusion zone is smaller and the macroscopic density is larger (see Table 1). Comparing the premigrational and postmigrational BCBP DRP, it is seen that the postmigrational dip is smaller. This is essentially a consequence of the mechanistic model: BCs attract BCBPs, thereby bringing some BCBPs laterally closer to other BCBPs.

The BC × BCBP cross DRP shows that before BCBP migration, the two populations are uncorrelated (shown by the flat DRP) as expected, but after BCBP migration the cross DRP is significantly non-flat and therefore the two populations are correlated. In particular, the postmigrational cross DRP has the shape of a Mexican hat. BCs have attracted some BCBPs from their immediate vicinity, thereby creating a higher than average density for small distances (*<* 20 μm) and lower than average density for distances up to around the maximal dendritic length of *l*_max_ = 44 μm. This non-flat profile is the fingerprint of the correlation between BCs and BCBPs.

Comparing the results of the Monte Carlo simulation (bars) with the real data (filled dots) shows that, in general, there is a good fit. In the case of the BC DRP and the BCBP DRP these results may be expected, given that the parameters of the model were chosen in order to secure a reasonable fit. However, the cross DRPs are very interesting. The real data falls quite nicely within the error bars of the simulated results and the shape of the simulated cross DRP closely follows the shape of the real data cross DRP.

For the other fields, the findings are qualitatively similar, both in the case of postmigrational cross DRP (compare Figure 10 and Figure 11), and for the other DRPs (data not shown). The mechanistic model is therefore capable of reproducing the observed correlation between BCs and BCBPs; this does not prove the model right, but it corroborates the underlying assumptions.

**Figure 11:**
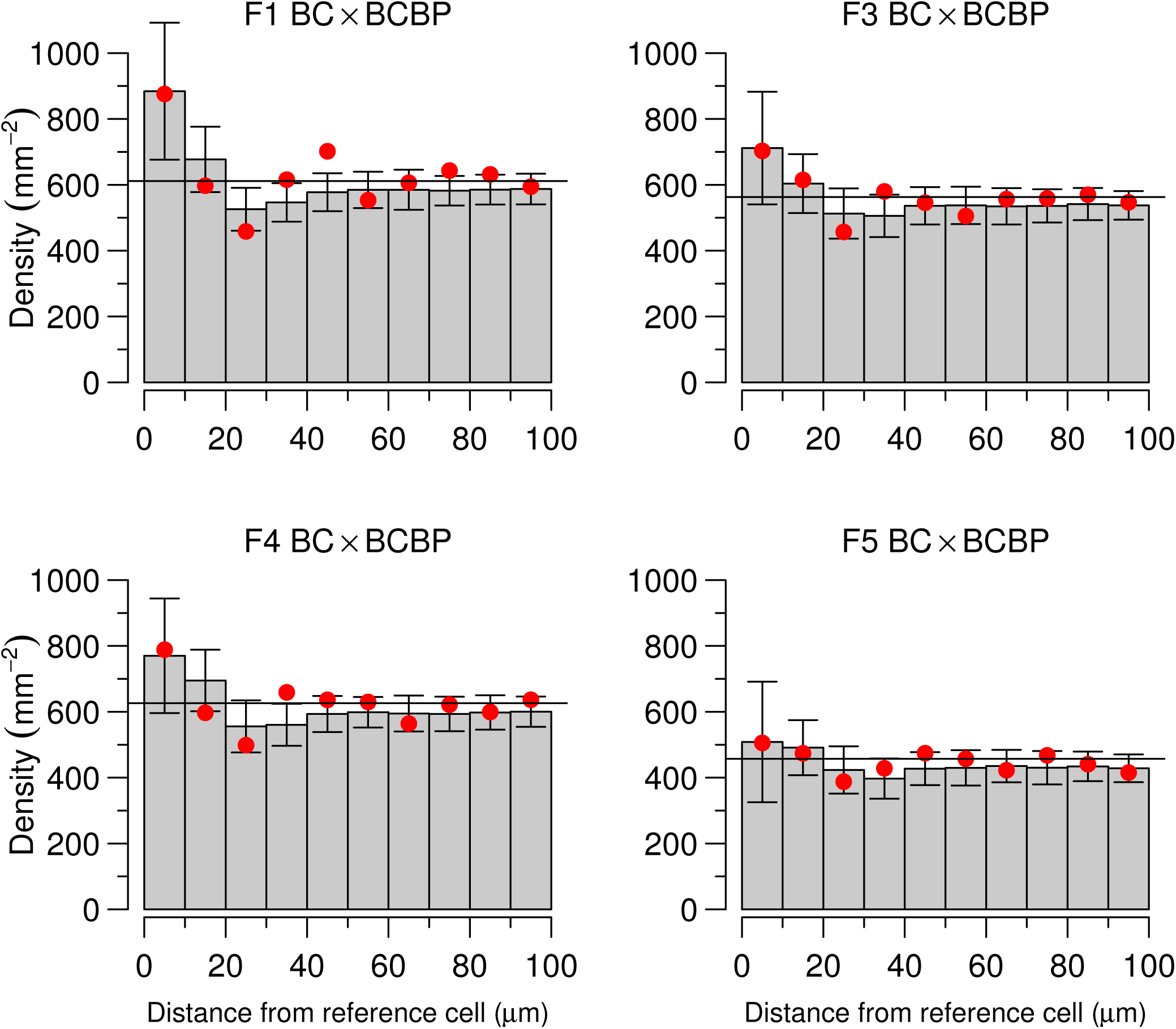
Postmigrational cross DRP results for fields 1, 3, 4 and 5. Cross DRPs are shown in the same format as in Figure 10.

Of course, these results depend on the choice of the relative friction-to-string force (see Table 2) and indeed the relative flat postmigrational cross DRP for field 5 (Figure 11) was the result of a mean relative force of *μ* _*f*/*s*_ = 0.97, somewhat larger than the mean relative forces for the other fields. Field 2 is the exception to this rule. Here, *μ* _*f*/*s*_ = 0.90, which is much higher than for field 1 (*μ* _*f*/*s*_ = 0.75) but despite this, the relative sizes of the peaks of those two fields are similar. Hence, field 2 is intrinsically more capable of attracting BCBPs to BCs and a higher friction force is needed to inhibit this mechanism. Within the framework of the model, this can probably be explained by the fact that the density of BCs and BCBPs are lower in field 2 and thus the BCBPs can migrate more freely towards the BC with less chance of colliding with other BCBPs (when simulating the model, collision of BCBPs was assumed to imply that migration stopped, as described in the methods, step 5).

## Discussion

A key reason for studying retinal mosaics is to reveal the developmental mechanisms that drive pattern formation. In this study, a mechanistic model for the interaction between BCs and BCBPs has been developed, simulated and assessed. The cornerstone of the model is that a BCBP can migrate due to a compromise between a string-like force exerted by the dendrite, connecting the BCBP with a BC, and friction forces within the INL. Using DRP analysis, it has been showed that the mechanistic model is capable of reproducing the empirically observed pattern of BC-BCBP interactions, as well as accounting for the properties of the homotypic mosaics and the number of synapses formed. Thus we propose our model as a candidate for the mechanism that generates the BC-BCBP correlation.

Other candidate models have previously been described. It has been suggested that a BC and a BCBP are daughters from a common progenitor cell (Chiu & Nathans, 1994), i.e. they are born at the same place and migrate radially. This hypothesis, however, fits poorly with the DRP results which shows that the correlation is far from perfect. At least this hypothesis should be supplemented by some explanation of why the BCs and BCBPs are not perfectly correlated. This explanation could be random fluctuations, but it is hardly satisfactory to explain a mechanism by referring to randomness as a key driver. Also, this model would probably not produce a Mexican hat shaped DRP but instead simply a DRP with a density that decreases monotonically with increasing distance. Finally, as pointed out by Kouyama & Marshak (1997), the fact that some BCBPs make two dendritic contacts seems to be at odds with the idea that they should connect only to sister BCs. Another candidate model was that of cell death, although this was already rejected as needing far too much cell death to generate the observed patterns (Kouyama & Marshak, 1997).

These considerations led Kouyama & Marshak (1997) to favour an explanation based on dendritic interactions and relative migration of BCs and BCBPs. They did not, however, develop an explicit model of this interaction. This gap has now been filled.

Another interesting empirical observation, which fits into the framework of the model, is that in marmoset (a New World monkey) BCs and BCBPs are positioned randomly in the ONL and INL, respectively, but conditionally on the positions of BCs, the BCBPs are nonrandom (Luo et al., 1999). In other words, BCs and BCBPs are also correlated in marmoset, but this correlation does not rely on either BC or BCBP neurons being arranged in regular mosaics. This is in accordance with the model developed here: the BCs are ‘leaders’ and via dendritic interactions with BCBPs, they induce BCBPs to migrate, i.e. to become ‘followers’, and this mechanism does not depend on the cells forming homotypic mosaics.

This notion of ‘leaders’ and ‘followers’ is supported by the observation that bipolar cells differentiate after cone photoreceptors (Rapaport, 2006). Hence, at the time of presumptive BCBP migration, the BCBPs may be more flexible and mobile than BCs, which is exactly what has been assumed in this study. BCBPs could therefore be thought of as ‘add-ons’, which adjust to the already established BC array.

In the framework of the model developed here, the migration of BCBPs can conceptually be decomposed into two parts: (i) migration caused by dendritic interactions with BCs and (ii) migration before a dendritic connection to a BC is made. A sanity check on the model developed here, is that it should not predict migration distances above 100 μm. (Estimates of migration distances for some cell types, but not BCBPs, suggest lateral movement to be well under 100 μm (Reese et al., 1999).) According to the output from the model, migration caused by dendritic interactions is only around 15 μm.

Based on these considerations, we suggest that the model developed in this study should be considered in future experimental and theoretical work on the developing retina. Future theoretical work might include conducting a sensitivity analysis of the parameters of the model, particularly of the relative friction-to-string force. Information about the sensitivity of the output with respect to the relative force would be valuable for future experimental work.

On the experimental side, techniques for measuring cellular forces are now available (such as optical tweezers) and measurements of dendritic string forces and retinal friction forces would provide a test of the model. In general, future experiments could provide tests of the assumptions made in this model. With a mechanistic model, testable hypotheses emerge much more naturally than in the case of a purely statistical model, and this is a strong argument for mechanistic models. Some assumptions could prove wrong and should be discarded, some might need an adjustment or more detail and some might be perfectly supported by experiments. In this manner, future experiments could spur new theoretical ideas which could then be built into future mechanistic models of the BC-BCBP interaction.

## Acknowledgements

The authors would like to thank Prof. Nobuo Kouyama (Tokyo Women’s Medical College) and Prof. David W. Marshak (University of Texas) for providing the data. The study was supported by a studentship from EPSRC (to AVL).

## 1 Postmigrational blue cone bipolar position

The cornerstone of the mechanistic model is the migration caused by a compromise between a dendritic string force and friction in the INL. In the article, the formula describing the postmigrational BCBP position was merely stated without derivation. This section presents the derivation.

BCBP migration is a linear motion and parallel with the INL. Thus, we only need to consider motion in one dimension. The physical situation is sketched in Figure 2 of the article.

Let **F**_*s*_ denote the dendritic string force exerted on the BCBP. Assume that the BC pedicle is immobile^1^ and that the BCBP can only move in a planar region and that the distance between the BC pedicle and this plane is *d*_*v*_. In this plane, orient an axis starting from the projection of the BC pedicle and pointing in the direction of the BCBP. Let *l* denote the position of the BCBP on this axis, i.e. initially *l* is positive and corresponds to the lateral distance between the BC pedicle and the BCBP. Let *θ* denote the angle which the connecting line between the BCBP and the BC pedicle forms with the *l* axis. Then the projection 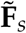 on the BCBP plane of the dendritic string force **F**_*s*_ has length 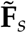 given by.

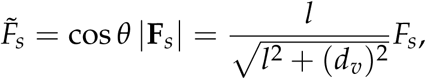

where *F*_*s*_ = *|***F**_*s*_*|*.

Within the INL, the BCBP cannot move freely, it is subject to a friction force. It is assumed that the static and dynamic friction forces are identical and have magnitude *F*_*f*_. They are obviously directed in the opposite direction of the initial 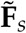. Motion requires that initially 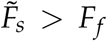, so this is assumed in the following derivations. This assumption is immaterial, since the case of no motion is trivial.

Given this geometry, the equation of motion of the system reads

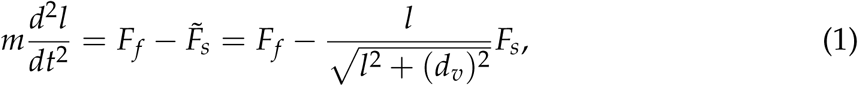

where *m* is the mass of the BCBP^2^. From this expression it is clear that it is only valid as long as the BCBP migrates in the direction of decreasing *l*, in other words if the BCBP changes direction at some time *t′*, then the expression is no longer valid because the friction force changes sign (it can be made valid again by simply reversing the *l* axis or reversing the sign of the friction force). In the following derivations, we shall assume that the BCBP will not change direction, i.e. no oscillations. We shall deal with oscillations below.

If we let *v* denote the velocity, the second derivative on the left hand side of this equation can be rewritten as

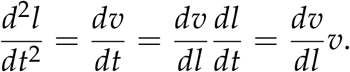

Note that it has been assumed that *l*(*t*) can be inverted such that we can express *v* as a function of *l*. This requires monotonic motion, but this is exactly what has been assumed above by ruling out oscillations.

Dividing Eq. (1) by *m* and using the last expression we obtain

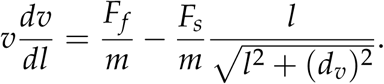

In this expression it is possible to separate the variables *v* and *l*. Hence,

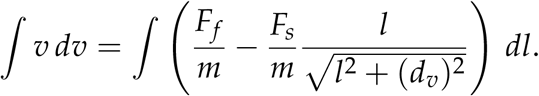

Performing the integration yields

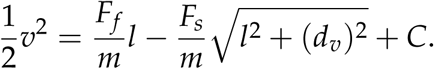

In order to determine *C* we can use that at *t* = 0, *l* = *l*_0_ (initial position) and *v* = 0. Inserting this into the above equation yields

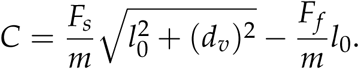

The equation governing the relation between position *l* and velocity *v* is therefore

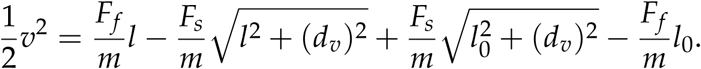

Now, we wish to determine the positions *l*^*\**^ where *v* = 0. We know that initially there will be motion (we assumed 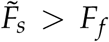). Because of the presence of friction, this motion cannot continue forever, eventually it will slow down and make *v* = 0, which means that either the BCBP has come to a stop or it is merely changing direction of motion. The latter can only be the case if *l*^*\**^ *<* 0, since only in this case can there be a force which will increase *l*. In this case, there will be oscillations. Because of the friction force, however, such oscillations will necessarily be damped, and therefore the BCBP will come to a stop at some point. If there is only one *l*^*\**^, except for *l*_0_, for which *v* = 0 and if this *l*″*\** is positive, then we know that 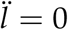 and oscillations will not be present. The BCBP will simply come to a stop at this *l*^*\**^ *>* 0. We will now show, that indeed there is only one such *l*^*\**^ and that this *l*^*\**^ is positive for realistic parameter values.

Imposing *v* = 0 in Eq. (1) and rearranging terms yields

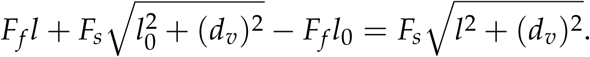

This can be rewritten as a quadratic equation in *l*, and therefore it can have a maximum of two solutions. The quadratic equation reads

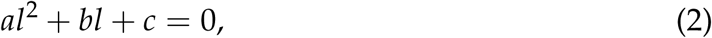

where

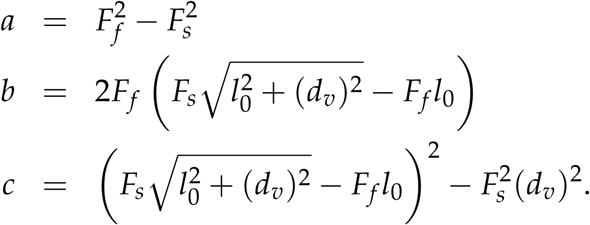

Note that *a ≠* 0, again because of the assumption that initially (at *l*_0_) 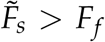, hence *F*_*s*_ *> F*_*f*_. Thus, Eq. (2) will never collapse into a linear equation.

After some tedious algebra, the discriminant of the quadratic Eq. (2) can be written as

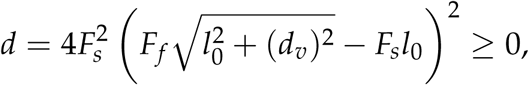

which is seen to be non-negative. It is only 0 in the case of 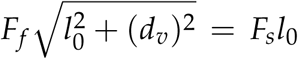, in which case there will be no motion. But this is assumed not to be the case, again by the assumption 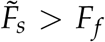 Hence *d >* 0 and there are two different real solutions, of which *l*^*\**^ = *l*_0_ must be one. The solutions are

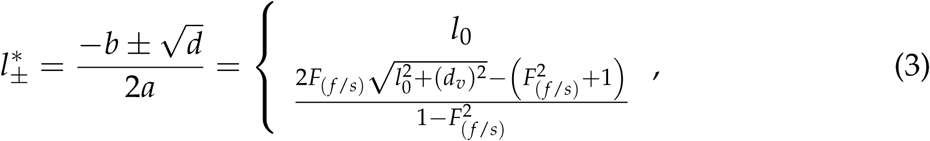

where we have introduced the shorter notation *F*_(*f*/*s*)_ = *F*_*f*_ /*F*_*s*_. Note that the denominator is always positive. Also note that the first solution 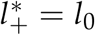 is trivial.

Finally, we need to argue that for plausible parameter values, it holds that 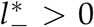, such that we can be sure that this 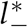 is not merely a point, where the BCBP changes direction (see above).

Hence, we need to know the sign of 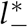. Based on the expression for 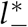 in Eq. (3) it follows that

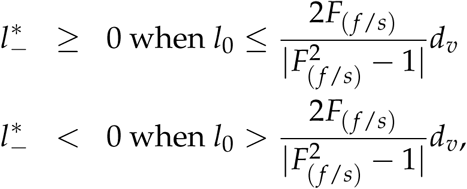

where it has been used that *F*_(*f*/*s*)_ *≠* 1.

For the values of *F*_(*f*/*s*)_ and *d*_*v*_ used in this study, it will very rarely happen that 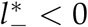 (e.g. five cases in an arbitrarily selected single simulation of all 1202 BCBPs from the five fields, i.e. 0.4 % of BCBPs.). For example, for field 2, *F*_(*f*/*s*)_ = 0.9, which implies that 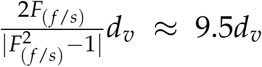. In turn, *d*_*v*_ is almost certainly larger than 10 μm. Lateral dendritic lengths *l* are never larger than *l*_max_ = 44 μm (see description of Step 4 in main article). We can therefore safely ignore the case of BCBP migration beyond the BC projection. Thus, in almost all cases, 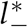 in Eq. (3) is indeed a point, where the BCBP comes to a stop.

## 2 Calculation of mean premigrational dendritic length

Prior to BCBP migration, BCs and BCBPs are spatially independent. We now consider a BCBP known to be migrating and thus known to be located within *l*_max_ of a given BC. Assuming that we know that this BCBP connects to this particular BC, the expected premigrational dendritic length is given by 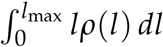, where *ρ*(*l*) is the density of BCBPs at a distance *l* from the BC. Because of the spatial independence between BCs and BCBPs, *ρ*(*l*) = *ql*^2^. It also holds that 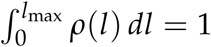. We can therefore calculate *q*

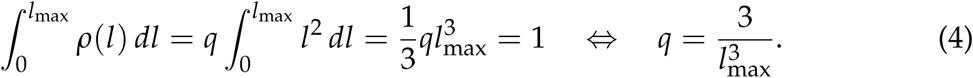

The expectation of *l* can now be expressed in terms of *l*_max_.

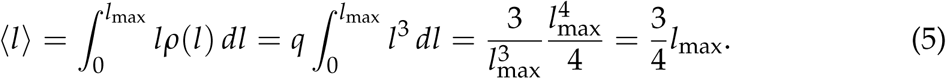

In our case *l*_max_ = 44 μm, implying that 〈*l*〉 = 33 μm. This analysis does not, how-ever, take into account that the larger *l* the smaller is the probability *ϕ*(*l*) that the BCBP will actually connect to the BC in question. This probability function *ϕ*(*l*) is much harder to estimate, but it explains why ϕ*l*〉 ≈ 20 μm instead of 33 μm.

## Footnotes

1 If the BC pedicle is also allowed to move, the force exerted on the BCBP will not only depend on BCBP position but also on BC position, and vice versa. Hence, the equations of motion would become coupled differential equations, which it is nice to avoid. Immobility of the BC pedicle is supported by biological observations, see article (methods, step 1) for details.

2 From this expression it is seen why a velocity dependent friction force is not appropriate. Assume that the friction force had the form *F*_*f*_ = *a* + *bv*. Solving the system requires us to find *l* such that *v* = 0 and *l* = 0. This implies that 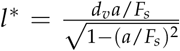, i.e. postmigrational dendritic length *l\** is independent of premigrationel dendritic length *l*_0_.

